# Using double cut *in vitro* assembled CRISPR/Cas9 to modify the genome of *Coccidioides posadasii*

**DOI:** 10.1101/2023.03.04.531116

**Authors:** Heather L. Mead, Daniel R. Kollath, Ashley N. Itogawa, Austin V. Blackmon, Matthew M. Morales, Mitchell L. Bryant, Marcus de Melo Teixeira, Bridget M. Barker

## Abstract

Although the genus *Coccidioides* is divided into two closely related and putatively allopatric species, analysis shows that hybridization has occurred between species and at least one *C. posadasii* conserved fragment has introgressed into several *C. immitis* genomes in a population-specific manner. Transcript abundance *in vitro* and *in vivo* for ten ORFs in this introgressed region were measured for several isolates. We used signals of introgression and high mRNA transcript levels in the spherule as indicators of selection for genes related to critical biological processes involved in *Coccidioides* pathogenesis. The only transcript in the introgression region with significant expression was a gene that encodes for a betadefensin-like (DEFBL) peptide rich in serines and cysteines. Few virulence factors have been identified in *Coccidioides*, and we employed the CRISPR-Cas9 mediated gene deletion tool to delete this gene in *Coccidioides*.

## INTRODUCTION

The genus *Coccidioides* comprises two species of pathogen, *Coccidioides immitis* and *C. posadasii*, which are the causative agents of the fungal disease coccidioidomycosis or “Valley fever.” These fungi are endemic to the southwestern United States, the Central Valley of California, arid regions of Mexico, Central and South America, with recent range expansions to areas such as Eastern Washington State (1). These thermal dimorphic fungi have two distinct life cycles, the saprobic where it grows as vegetative mycelia in the soil and the conversion in the host to an endospore filled spherule (2). This dramatic shift in morphology is a key virulence factor for *Coccidioides* (3). Understanding the mechanisms behind this switch is key to unraveling pathogenesis, the evolution of virulence, and potential drug targets.

However, to assess function, researchers must have a tool kit of molecular methods, such as the ability to delete, modify, and insert genes. To date, there are few molecular tools and a relatively small number of gene knockouts in *Coccidioides* (4, 5, 6, 7). This lack of tools and detailed knowledge of the function of proteins regulating the morphological switch limits understanding of virulence in *Coccidioides*. Strains of the fungus that have lost the ability to make intact spherules have a significant decrease in the virulence phenotype *in vivo* (3, 4, 8). Therefore, defining the genes involved in this morphological shift is crucial to understanding virulence.

One method to survey the genome for targets is to complete RNAseq experiments. Several transcriptomic studies have examined differences in gene expression in the mycelia (saprobic stage) compared to the spherule (parasitic stage) to identify genes critical to morphogenesis (3, 9, 10, 11, 12). In Whiston *et al*. 2012, a significant number of genes were found to differ between the two life stages, namely cell surface-associated genes, particularly chitin-related genes as well as potential virulence factors such as alpha (1,3) glucan synthase, SOWgp, and several genes in the urease pathway were upregulated in the parasitic life stage. A similar study compared transcripts between the life stages as well as wild type to an attenuated strain of *C. posadasii* that does not form intact spherules (12). The results of this study showed 252 genes that were conserved and improperly expressed in the mutant stain. One gene of interest that was shown to have an improper response in the mutant strain was predicted to be a heat shock protein that may be expressed during environmental stressors such as entering a host body. These studies clearly show differences in gene expression between the two life stages; however a functional assessment of these hypotheses need to be completed.

To definitively show that these genes are important for virulence, mutant strains must be created. This has been a challenge in *Coccidioides*. Traditional genetic approaches are either inefficient or nonfunctional in filamentous fungi due to multiple nuclei per conidium (heterokaryotic cells) and low homologous recombination efficiency (13). The challenges of genetic manipulation in non-model filamentous fungi, such as *Coccidioides* spp., make methods such as in vitro assembled CRISPR/Cas9 a very powerful tool to examine gene function (14). Extensive CRISPR/Cas9 systems have been successfully developed for model fungi, such as *S. cerevisiae* (15), but more recently have been applied in non-model filamentous fungi such as *Blastomyces dermatitidis*, a close relative of *Coccidioides* (16). In *B. dermatitidis*, CRISPR/Cas9 was used to disrupt genes involved in zinc uptake to determine that this mechanism was important upon infection of a host (17). Despite being a medically important pathogen, no successful CRISPR/Cas9 tools have been reported for *Coccidioides* to date.

To remedy this deficit, we targeted CIMG_00509/CPSG_05265, which has been identified in studies as being highly upregulated in the parasitic growth phase compared and associated with a region of introgression (3, 10, 18). It was first identified at a locus of introgression between the two species containing several genes, from CIMG_00498 to CIMG_00509 in the *C. immitis* genome of strains RS (18). Upon later completion of RNAseq, this region showed low levels of expression in both life stages, except for CIMG_00509, suggesting that this gene is involved in the development of spherules. This evidence points to CIMG_00509 being an important gene upon the infection of a host and therefore an ideal candidate to delete. Several attempts were made with molecular techniques used in *Coccidioides posadasii* to delete the orthologous locus CPSG_05265, but all were unsuccessful (19, 20). This led us to develop an *in vitro* assembled double-cut CRISPR/Cas9 methodology to delete CIMG_00509. The result is the first successful application of CRISPR/Cas9 technology in *Coccidioides*.

## METHODS

*Strains* The wildtype strain Silveira (NR-48944) was used for trasnformation (21). The plasmid pCB1004 from the fungal genetics stock center was used for the template of the hph gene.

### Creation of selection marker repair template

Hi-fidelity PCR was completed with ExTaq and 100mM primers: P1_HYG_fwd,CGACGTTAACTGATATTGAAGGAGCATTTTTTGGGCTTGGC; P2_HYG_rev,GTTAACTGGTTCCCGGTCGGCATCTACTCTATTCCTTTGCC; P3_HYG_509fwd,CTTTAGGGCAACCATGTAATCCAAGCTCTGACGACGTTAACTGATATTGAAGGAGCATTTTTTGGGCTTGGC; P4_HYG_509rev,GTTAACTGGTTCCCGGTCGGCATCTACTCTATTCCTTTGCCGCGAAGAAGCCTTGTGCCTGTGATCAGCAAC; and the hph template to create a ∼1.4 kb linear DNA repair template using an overlap PCR protocol similar to (22).

### Creation and transformation of protoplasts

Protoplasts were created as described (20, 23). Briefly, 100 ml of liquid 2xGYE was inoculated with 5 × 10^8^ *C. posadasii* strain Silveira arthroconidia and on a shaking incubator at 30°C for ∼17h until germlings were observed by light microscopy.

Germlings were centrifuged at 2,800x g for 10min at 10°C, media was replaced with cell wall digestion buffer (chitinase 1.17mg/ml, drislase 7.58mg/ml, and lysing enzymes 4mg/ml) in osmotic buffer A (50mM potassium citrate, 0.6M KCl pH 5.8) and incubated on a shaking platform at 30°C for 75 min. Protoplasts were pelleted by centrifugation at 900 × g for 10 min at 10°C and resuspended in osmotic buffer B (10mM sodium phosphate, 1.2M MgSO_4_, pH 5.8). The trapping buffer (100mM MOPS, 0.6M sorbitol, pH 7.5) was gently added to create a density gradient layer and protoplasts were centrifuged using a swinging bucket at 5,000 RPM for 15 min at 10°C. Protoplasts were removed from the interphase layer and placed on ice in the MOPS buffer containing sorbitol (MS buffer: 10mM MOPS, 1M sorbitol pH 6.5). Finally, cells were washed twice in MSC buffer (MS buffer with 20mM CaCl_2_).

### DNA integration and chemical selection

Detailed protocol is available (24). Approximately 1×10^7^ protoplasts were incubated with 2.4ug of template DNA, Cas9:crRNA:tracrRNA ribonucleoprotein (RNP) complex and 8x the volume of 60%PEG solution on ice for 50 minutes, the reaction concluded with the addition of 3x the volume 60%PEG. Cells were combined with pre-warmed GYES soft agar and overlaid onto a pre-warmed GYES agar in plate. Protoplast s were allowed to grow undisturbed for 48hr at 30°C. Next GYE soft agar with hygromycin (75 μg/mL) was overlaid on the colonies and incubated in the dark for 4 days at 30°C. Single colonies were selected from plates and transferred to fresh 2xGYE hygromycin (75 μg/mL) plates. Colonies were replated every 5-7 days, several potential successful colonies were identified and subjected to gDNA extraction using phenol chloroform as described below.

### Mutant Screening

Genomic DNA was amplified using primers designed to amplify i) hygromycin repair template, ii) the native wildtype locus, iii) upstream flanking region + mutant locus, iv) downstream flanking region + mutant locus (Table 1). Four potential mutants were screened for all target regions, and wildtype C. posadasii strain Silveira was used as a control. Putative mutants were initially screened using PCR for presence of the *HPH* selection marker (Hygromycin B Resistance cassette) and absence of the wildtype gene sequence. Additional primers including regions upstream and downstream of the knockout locus were also included.

**Table 1.**
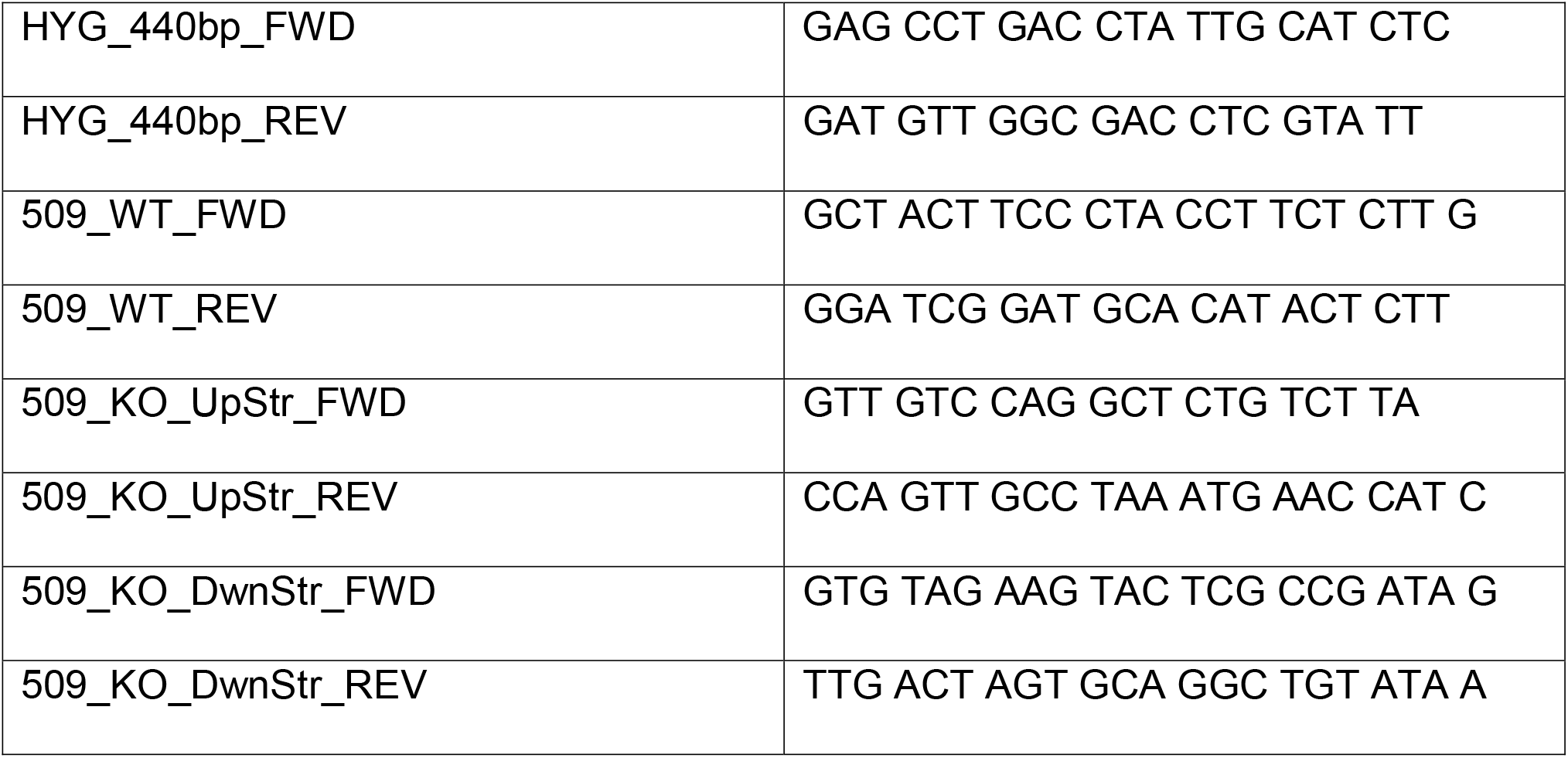
Primer sequences used to screen putative transformants.

### DNA extractions

Genomic DNA was extracted via phenol chloroform DNA extraction and ethanol precipitation after heat kill of mycelia in the BSL3 laboratory (25). This was paired with liquid nitrogen grinding to ensure efficient cell lysis to increase DNA yield. This approach generally resulted in sufficient quality and quantity DNA for PCR, as determined by gel electrophoresis and Qubit (ThermoFisher) quantification methods as previously described (26). For Southern blot analysis, higher quality DNA was required. A detailed protocol for high molecular weight (HMW) DNA extractions is available at (27).

### Southern Blots

A Southern blot was performed on putative mutants to further confirm removal of the target gene using protocol from (28). First, 3ug of HMW genomic DNA from both mutants and strain Silveira was digested with PvuI. All strains were run on a gel and blotted to a positively charged membrane via vacuum blotting. The membrane was hybridized with a DIG-labeled PCR-generated probe, complementary to a 616bp sequence upstream of the 509/5265 locus. The membrane was then washed and detected using the Roche DIG Detection Kit and imaged using a UVP protein imager. The fragment complementary to the DIG-labeled probe is predicted to hybridize to a 13,463bp digest fragment in Silveira and to a 4,074bp fragment in a successful transformant.

## RESULTS AND DISCUSSION

### Analysis of transcript differences between the two life phases

of *Coccidioides* spp. revealed the abundance of the transcript was 24-fold higher in the parasitic phase than the saprobic phase (10). Our initial investigation of the using RT-PCR of DEFBL *in vivo* from total RNA extracted from mouse lungs indicates it is also highly expressed during infection (data not shown). However, the function of the gene is elusive, with greatest homology to a beta-defensin-like peptide in *Anole carolinensis* and toxins in spider venom using blastN search on NCBI. Only one minor homologous hit in a fungal organism was found in the entomopathogenic bio-control agent *Metarhizium anisopliae*. Analysis using SignalP 4.0 suggests the protein is cleaved and secreted (29). Classic betadefensins in vertebrates are characterized by six cysteine residues in the form C-X6-C-X4-C-X9-C-X6-C-C. The sequence motif in both species of *Coccidioides* is C-X6-C-X5-C-X7-C-X5-C-X-C; thus, we are using beta-defensin-like peptide as our current nomenclature (30). DEFBL genes commonly encode toxic peptides of reptiles and anti-microbial peptides in various organisms. These small secreted peptides typically consist of a signal peptide, a pro-segment sequence followed by a ∼30AA defensin with the six cysteine residues that create 3 disulfide bridges and are 2-6kD. Sequence analysis confirms that DEFBL in both species of *Coccidioides* contain all these features. Protein alignment also indicates that there are 3 amino acids deleted in the *C. immitis* sequence type, and an arginine to leucine change at position 71. Interestingly, the *C. posadasii* sequence type of the gene is found in *C. immitis* strain RS, which is one of the common virulent lab strains (18).

We created a gene deletion in *C. posadasii* using the double-cut CRISPR-Cas9 technology with success. We have confirmed deletion of DEFPL-cp using PCR and Southern blot. No obvious defects in growth rate or production of spherules were evident. We have used the technique to target other genes and suggest that 100bp of homology results in better transformation efficiency and using synthetic DNA rather that the more laborious overlap PCR is an improvement that should be explored.

## SUMMARY

In summary, we demonstrated that the CRISPR/Cas9 tools can be applied to *Coccidioides posadasii* comparable to results in other fungi (Kujoth et al. 2018). The CRISPR/Cas9 system worked well for this difficult target compared to previously described gene deletion methods in *Coccidioides*. This is a substantial gain in the ability to understand gene function in *Coccidioides* which can lead to better understanding of virulence factors.

**Figure 1.**
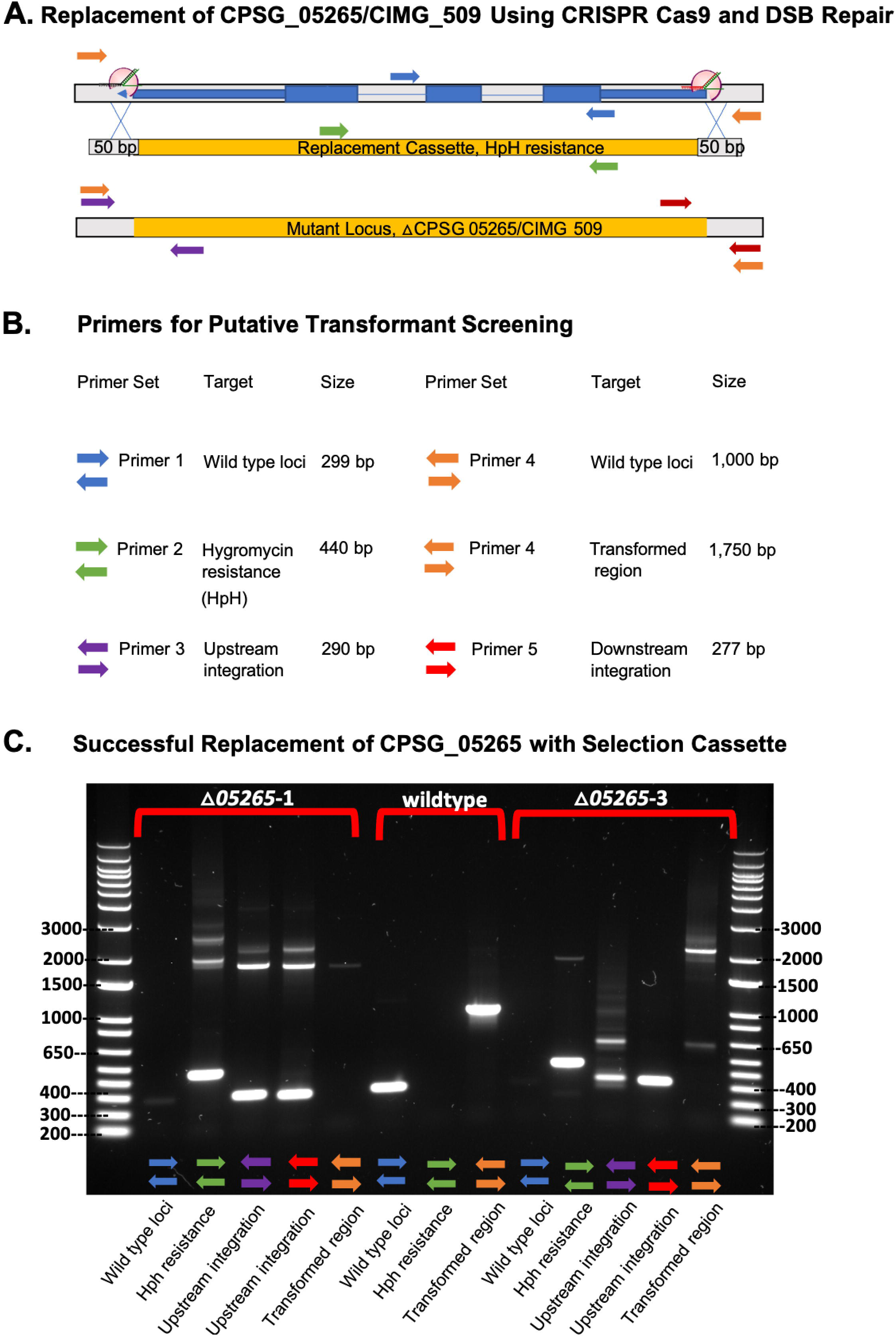
A. Schematic of CRISPR/Cas9 construct to replace CPSG_05265. B. Table of primers and predicted PCR results. C. Gel image of 2 transformants (5-1 and 5-3) and wildtype (middle lane) PCR results. Both transformants are hyg+ (green primer set) and show loss of target (blue primer set). Amplifying across the whole locus (orange primer set) indicates that 5-3 has integrated the repair construct and Southern blot shows complete loss of wildtype nuclei for this strain.

**Figure 2.**
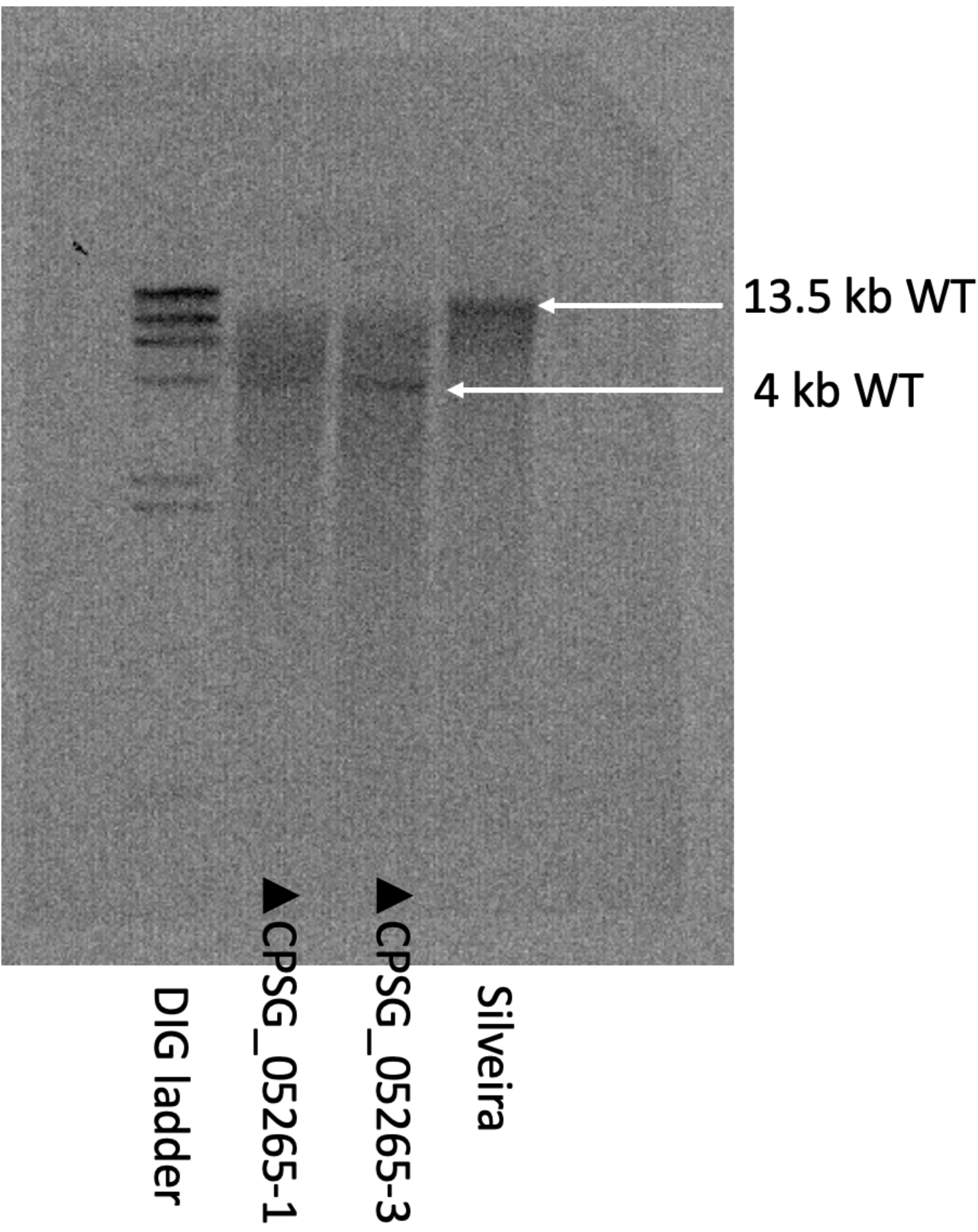
Southern blot of DNA extracted from 2 transformants in Figure 1 probed with 616bp DIG-labeled probe to produce a 13,463bp digest fragment in the wildtype strain and a 4,074bp fragment in a successful transformant. Lane 1 is the Roche DIG ladder, Lane to is transformant 5-1

